# Altered Dynamics of Cortical Beta-Oscillations during Motor Learning in Cerebellar Ataxia

**DOI:** 10.1101/2020.10.06.328807

**Authors:** Jana Klimpke, Dorothea Henkel, Hans-Jochen Heinze, Max-Philipp Stenner

## Abstract

Cerebellar ataxia is associated with an implicit motor learning dysfunction, specifically, a miscalibration of internal models relating motor commands to state changes of the body. Explicit cognitive strategies could compensate for deficits in implicit calibration. Surprisingly, however, patients with cerebellar ataxia use insufficient strategies compared to healthy controls. We report a candidate physiological phenomenon of disrupted strategy use in cerebellar ataxia, reflected in an interaction of implicit and explicit learning effects on cortical beta oscillations. We recorded electroencephalography in patients with cerebellar ataxia (n=18), age-matched healthy controls (n=19), and young, healthy individuals (n=34) during a visuomotor rotation paradigm in which an aiming strategy was either explicitly instructed, or had to be discovered through learning. In young, healthy individuals, learning a strategy, but not implicit learning from sensory prediction error alone, decreased the post-movement beta rebound. Disrupted learning from sensory prediction error in patients, on the other hand, unmasked effects of explicit and implicit control that are normally balanced. Specifically, the post-movement beta rebound increased during strategy use when implicit learning was disrupted, i.e., in patients, but not controls. We conclude that a network disturbance due to cerebellar degeneration surfaces in imbalanced cortical beta oscillations normally involved in strategy learning.

## Introduction

Accurate arm movements are fundamental to everyday human behavior, e.g., when reaching out to shift gears while steering a car. Systematically inaccurate reaching, such as in cerebellar ataxia, drastically impairs quality of life. Inaccurate reaching in cerebellar ataxia has been explained by an implicit motor learning dysfunction, specifically, by a miscalibration of internal models that relate outgoing motor commands to resulting state changes of the body and its environment (Smith and Shadmehr, 2005; Tseng et al., 2007; Synofzik et al., 2008; Izawa et al., 2012; Bhanpuri et al., 2014). In the motor learning literature, cerebellar ataxia is therefore often viewed as a prototypic disorder of motor adaptation, which has long been regarded primarily as a process of recalibrating an internal model (e.g., Palmer, Auksztulewicz, Ondobaka, & Kilner, 2019; Tan, Wade, & Brown, 2016; Thoroughman & Shadmehr, 2000).

In the last decade, however, motor adaptation has been increasingly recognised as the result of several interacting learning mechanisms (Huang et al., 2011; Izawa and Shadmehr, 2011; Codol et al., 2018; Kim et al., 2019). These include learning cognitive strategies, e.g., learning to explicitly re-aim when reaching is inaccurate (McDougle, Ivry, & Taylor, 2016; McDougle & Taylor, 2019; Taylor & Ivry, 2011; Taylor, Krakauer, & Ivry, 2014). In theory, such alternative learning mechanisms could help patients with cerebellar ataxia compensate for miscalibrated internal models (Therrien et al., 2016; Donchin and Timmann, 2019). However, cerebellar pathology seems to cause an additional disruption in developing, using, or maintaining cognitive strategies for aiming (Vaca-Palomares et al., 2013; Butcher et al., 2017). Recently, Wong et al. (2019) proposed that this disruption arises from an interaction between strategy learning and internal model adaptation, specifically, from a suppression of strategy learning by internal model adaptation, even when the latter is impaired. What could be the physiology underlying this interaction, and its pathophysiology in cerebellar ataxia?

A physiological signal that has recently attracted considerable attention as a candidate signal involved in motor adaptation is the so-called post-movement beta rebound (Alayrangues, Torrecillos, Jahani, & Malfait, 2019; Palmer et al., 2019; Palmer, Zapparoli, & Kilner, 2016; Tan et al., 2016; Tan, Jenkinson, & Brown, 2014; Torrecillos, Alayrangues, Kilavik, & Malfait, 2015). Movement is accompanied by characteristic peri-movement modulations of beta-power in cortical and subcortical sensorimotor regions (van Wijk et al., 2012), which can be recorded using electroencephalography (EEG). Immediately before and during movement, the amplitude of sensorimotor beta-oscillations decreases, followed by an increase above baseline, called the post-movement beta rebound (Pfurtscheller, 1992). Several studies have demonstrated that the sensorimotor cortical post-movement beta rebound decreases at early stages of visuomotor adaptation, as well as force-field adaptation, and increases again via later stages of adaptation (Alayrangues et al., 2019; Palmer et al., 2019; Tan et al., 2016; Tan et al., 2014; Torrecillos et al., 2015). While these reports are compatible with a framework in which beta-oscillations have a more general role in maintaining vs. releasing a “status quo” (Engel and Fries, 2010; Brittain and Brown, 2014; Hosaka et al., 2016), the exact learning process(es) reflected in the dynamics of the post-movement beta rebound are unknown. First and foremost, the design of previous studies has prevented any dissociation between internal model adaptation and strategy learning. This dissociation, however, is critical when asking how the two learning processes may interact physiologically, and how this interaction may differ in cerebellar ataxia. To date, learning-related dynamics of cortical signals in cerebellar ataxia have remained unstudied.

Here, based on a behavioral paradigm that isolates internal model adaptation from strategy learning, we examine learning-related dynamics of the post-movement beta rebound in cerebellar ataxia. We first determine the dynamics of the post-movement beta rebound when healthy individuals have to learn an aiming strategy for reaching. We then ask whether the same signal also depends on intact internal model adaptation, and how its dynamics differ in cerebellar ataxia, i.e., when internal model adaptation is disrupted. Our study thus extends a hitherto small, but growing literature that adopts a network perspective on the electrophysiology of cerebellar ataxia (Lu et al., 2008; Aoh et al., 2019; Visania et al., 2020).

To disentangle strategy learning and internal model adaptation, we employ a behavioral paradigm introduced by Mazzoni and Krakauer (2006). When healthy individuals are instructed an aiming strategy that fully compensates for a visuomotor rotation, e.g., by aiming 45° clockwise of the target when the rotation is 45° counter-clockwise, use of that strategy eliminates target error, at first. That is, the (rotated) visual representation of the hand initially moves in the desired direction, towards the target. Surprisingly, however, healthy individuals soon start overcompensating, i.e., moving in a direction that is further and further away from the strategic aim point and the target (>45° clockwise in the above example). This is because an aiming strategy does not eliminate the sensory prediction error that results from a discrepancy between the seen movement direction, and the direction that is implicitly expected based on an internal model of visuomotor contingencies. Sensory prediction error is the driving force for an implicit recalibration of that visuomotor model, a recalibration that gradually overrides the aiming strategy (Mazzoni & Krakauer, 2006; Taylor & Ivry, 2011). Importantly, patients with cerebellar ataxia overcompensate less in this task, and therefore perform better, on average (Taylor, Klemfuss, & Ivry, 2010), in line with the idea of a recalibration deficit in cerebellar ataxia.

By comparing two groups of young, healthy individuals, one of which deployed an instructed strategy, while the other had to discover a strategy, we isolate effects on the post-movement beta rebound related to strategy learning, over and above learning purely from sensory prediction error (Experiment 1). In Experiment 2, we then ask for effects of sensory prediction error, or learning from sensory prediction error, on the post-movement beta rebound, and for changes of this signal in cerebellar ataxia. We do so by comparing patients with cerebellar ataxia and healthy, age-matched controls when both use an instructed aiming strategy to counter a visuomotor rotation, effectively eliminating the need for strategy learning. Our results indicate that both explicit and implicit control share as a common neural pathway a modulation of the post-movement beta rebound, a modulation that is out of balance in cerebellar ataxia. We propose that this imbalance represents a candidate physiological phenomenon of impaired strategy learning in cerebellar ataxia.

## Materials and Methods

### Subjects

Thirty-four young, healthy, right-handed individuals participated in Experiment 1 (mean ± standard deviation: 26.3 ± 6.04 years; 20 female). Eighteen right-handed patients with cerebellar ataxia and nineteen right-handed, age-matched healthy participants took part in Experiment 2. Three patients were excluded from analyses because post-hoc interviews revealed that they had misunderstood task instructions (specifically, they misunderstood the compensatory strategy required in the behavioral task). Included patients (eight female; **Table 1**) were, on average, 57.5 ± 12.5 years old (mean ± standard deviation; healthy controls: 56.4 ± 10.3 years, seven female). Patients were recruited at the Department of Neurology at Otto-von-Guericke University and via national support groups. Symptom severity was assessed using the International Cooperative Ataxia Rating Scale (ICARS; Trouillas et al., 1997). Group sizes for both experiments were chosen to be at least as large as in previous EEG and behavioural studies into motor adaptation in healthy individuals and patients with cerebellar ataxia (Aoh et al., 2019; Tan, Wade, & Brown, 2016; Tan, Jenkinson, & Brown, 2014; Taylor, Klemfuss, & Ivry, 2010; Torrecillos, Alayrangues, Kilavik, & Malfait, 2015). All participants had normal or corrected-to-normal vision, and all gave written informed consent according to the Declaration of Helsinki. The study was approved by the local ethics committee at Otto-von-Guericke University Magdeburg.

### Experimental Setup

Experiments were conducted in a dimly lit, electrically shielded, and sound-attenuated chamber. Participants were sitting at a graphics tablet placed on a table in front of them (Intuos Art, Wacom, Kazo, Japan; 2540 lines per inch, sampled at 60 Hz, active area of 21.6 × 13.5 cm). They were facing an LCD monitor placed on the table at a distance of 70 cm (52 × 32.4 cm, refresh rate 60 Hz). During the experiment, participants moved a stylus held in their right hand across the graphics tablet while the monitor provided visual feedback of the current stylus location at 60 Hz (**Fig. 1A**). The tablet, stylus, and right hand were hidden from sight beneath a box.

**Figure 1.**
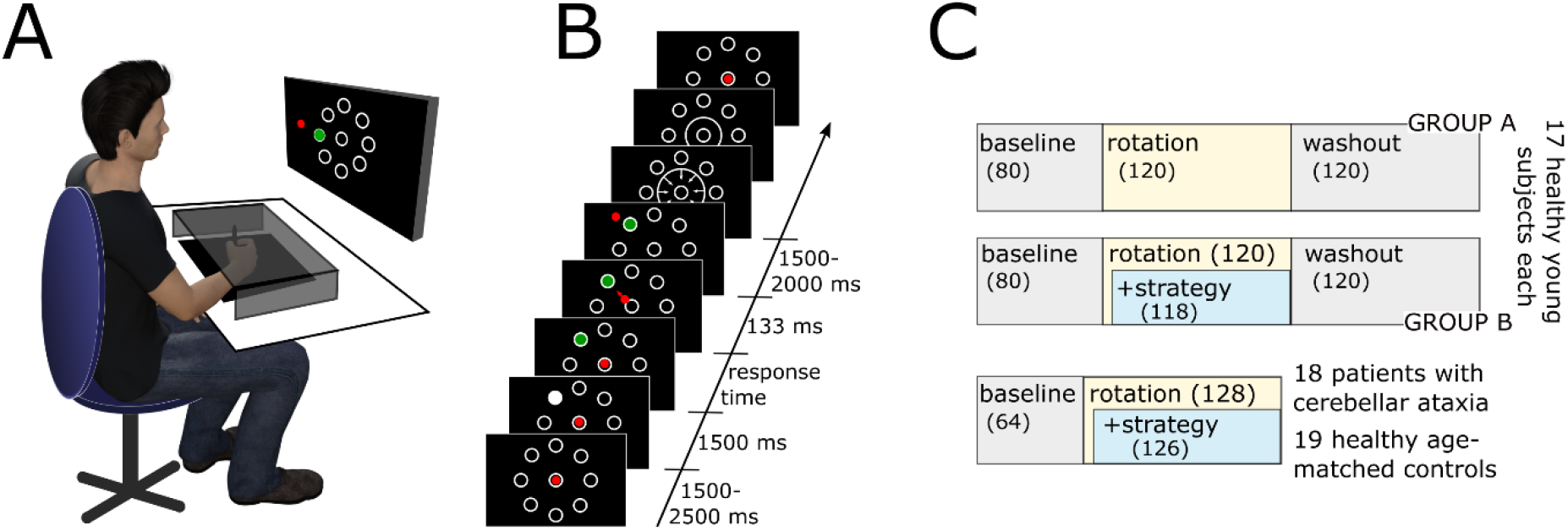
Experimental Setup (A), schematic of a single trial (B) and study design (C). In C, the upper two rows show the design of Experiment 1, and the bottom row shows the design in Experiment 2.

**Figure 2.**
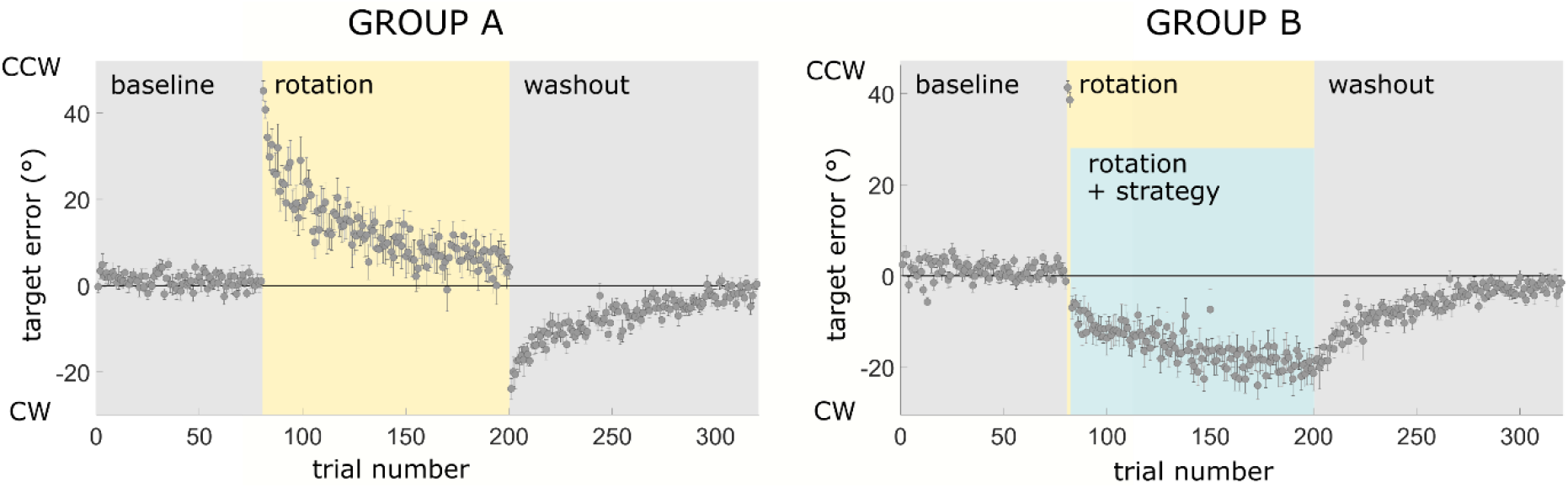
Learning in Experiment 1. Target error as a function of experimental block (baseline, rotation/rotation + strategy, washout) in group A (left) and B (right). Grey dots and error bars represent the mean and standard error of the mean, respectively, across participants in each group. CCW, counter-clockwise; CW, clockwise.

Experiments 1 and 2 consisted of a centre-out reaching task. Throughout both experiments, participants saw eight white, outlined target circles (radius 0.45° visual angle) arranged in a circular array (radius 6.04° visual angle, i.e., 7.4 cm) around the screen centre in steps of 45° (including cardinal directions; Fig. 1A). The screen background was black. These circles represented alternative target directions for reaching movements required in the task. At the centre of the screen, there was another white, outlined circle of the same size representing the required starting position (home position) for each reach. A solid red dot (radius 0.3° visual angle) on the screen represented current stylus location on the tablet (except at the end of each trial, see below). The red dot was at the centre of the home position when the stylus was at the centre of the tablet. To minimize movement artefacts in EEG, the task required stylus movements of only a few centimetres. Due to a technical issue, the spatial transformation of stylus movement on the tablet to movement of the red dot on the screen was anisotropic (see discussion). To move the red dot along the screen’s vertical axis from the home position to the circular array of targets, the stylus had to move from the tablet centre 1.01 cm along the tablet’s y-dimension (away or towards the participant). To move the red dot along the screen’s horizontal axis from the home position to the circular array of targets, the stylus had to move from the tablet centre 1.62 cm along the tablet’s x-dimension (parallel to the participant’s chest).

### Experiment 1

Experiment 1 asked whether, and how, the post-movement beta rebound changes when healthy individuals learn a strategy for reaching, over and above any changes when learning is purely driven by sensory prediction error. To address this question, we compared two groups of young, healthy individuals, each tested in a visuomotor rotation paradigm (n=17 participants in each group, age: 25 ± 5.4 years, and 27.5 ± 6.6 years, t(32)=1.2, p=.23). The two groups completed the same experimental paradigm except for one aspect of the instructions they received (**Fig. 1C**, top). The task for both groups was to “shoot” the red dot from the home position through one of the eight target circles with a single, rapid, straight movement, and to stop anywhere beyond the target. The target changed from trial to trial, and was highlighted at the beginning of each trial (see below). Both groups first completed a baseline block in which stylus movements along x- and y-dimensions of the tablet moved the red dot horizontally and vertically on the screen, respectively (e.g., a stylus movement forward, i.e., away from the participant’s body, and to the right moved the red dot on the screen upward and to the right). Following baseline, the red dot was rotated 45° counter-clockwise in both groups. Learning to compensate for this rotation involves both implicit adaptation of an internal visuomotor model, driven by sensory prediction error, and learning an explicit aiming strategy (Mazzoni and Krakauer, 2006), in this case, to aim further clockwise. To isolate effects of strategy learning over and above pure implicit learning, the two groups received different instructions once the visuomotor rotation was introduced. One group (group A) was not explicitly informed about the presence or nature of the visuomotor rotation, and received no instruction how to compensate for it. The other group (group B) was explicitly informed about the nature of the visuomotor rotation after the first two trials in which that rotation was present. Importantly, this group also received instructions to use an aiming strategy that could fully compensate for the visuomotor rotation, i.e., to aim at the neighbouring target in the clockwise direction. Since the imposed visuomotor rotation was identical in both groups, we assumed similar sensory prediction error-driven learning across groups, in line with Mazzoni & Krakauer (2006). Thus, any differences in post-movement beta rebound between groups were expected to reflect different demands on strategy learning, which was rendered unnecessary, by instruction, in group B, but not in group A (we include an alternative interpretation of group differences with respect to error in our discussion).

In group B, sensory prediction error-driven learning was expected to result in a gradual overcompensation (Mazzoni and Krakauer, 2006). To avoid any adjustments to the instructed strategy driven by the resulting systematic, gradually increasing target error (Taylor & Ivry, 2011), participants in group B were told to strictly adhere to the instructed strategy throughout the entire rotation block.

Across all blocks, and in both groups, the sequence of events in any given trial was as follows (**Fig. 1B**). First, the home position turned solid white and participants moved the red dot into the home position. After holding it in place (≤ 0.45° visual angle from the screen centre) for 1500-2500 ms (uniform random distribution), the home position turned back into an outlined, empty circle, and one of the eight target circles turned solid white instead. Participants were instructed to wait for that target to turn from solid white to solid green (after 1533 ms) before “shooting” the red dot through that target with a single, rapid, straight movement. The red dot “froze” in place between 117 and 133 ms after leaving the home position (depending on the time of movement initiation relative to the monitor’s refresh phase). Participants were instructed to move fast enough so that, by the time the red dot “froze” in place, it had passed the target. Upon stopping, participants had to hold the stylus in place for 1500-2000 ms (uniform random distribution). The red dot was then replaced by a white, outlined circle centred on the home position, whose radius decreased or increased, depending on the distance of the stylus from the centre of the tablet. This circle thus guided participants in their return to the home position without providing information about current direction relative to the home position. Thus, when the visuomotor rotation was present, participants could only learn about the rotation from centre-out movements, not from return movements. The red dot re-appeared and replaced the return circle upon reaching a radius of 1.55° visual angle around the screen centre.

After familiarising themselves with the experimental setup and task, both groups of participants first completed 80 baseline trials, followed by 120 rotation trials and 120 washout trials (in which the visuomotor mapping was identical to the baseline block). In each of these three blocks, each of the eight target locations was selected equally often, with the order of target locations pseudo-randomized across trials. Between the rotation and washout block, participants in group B were informed that the visuomotor rotation was switched off again, and asked to aim directly at the highlighted target, as in the baseline block. Participants in group A received no such instructions.

### Experiment 2

While Experiment 1 isolated effects of strategy learning, Experiment 2 asked for sensory prediction error-driven learning effects on the post-movement beta rebound, and for consequent changes of this signal in cerebellar ataxia. To this end, Experiment 2 compared patients with cerebellar ataxia, who learn less from sensory prediction error (Taylor et al., 2010), and healthy age-matched controls, in a paradigm that eliminates the need for strategy learning. This paradigm was identical to the one employed in Experiment 1, with few differences. First, both groups, i.e., patients as well as controls, received the same instructions, which were identical to the instructions given to group B in Experiment 1. That is, in both groups, strategy learning was rendered unnecessary by informing participants about the nature of the visuomotor rotation, and by instructing a fully compensatory aiming strategy. Second, several changes to the paradigm aimed at facilitating task performance for patients. Specifically, both groups were allowed to move more slowly than in Experiment 1 (the red dot “froze” in place 150 to 166 ms after leaving a radius of 1.6° visual angle around the home position). In addition, in both groups, the red cursor re-appeared earlier upon return than in Experiment 1 (at a radius of 3.15° visual angle around the centre of the screen). Also, the requirement to hold the stylus in place for the next trial to start was less strict (the red dot had to stay within a radius of 1.6° visual angle relative the centre of the screen). Finally, there were overall fewer trials (64 baseline trials, followed by 128 rotation trials, and no washout). Third, to ensure that outward reaching movements involved no, or only minimal, feedback corrections, the visuomotor rotation was unexpectedly switched off in 24 trials randomly distributed across the rotation block (catch trials). The rationale was that the curvature of movement trajectories would not differ significantly between these catch trials, and regular rotation trials, if feedback corrections had only minimal, if any, influence on movement trajectories (**Supplementary Material**). Apart from this comparison, catch trials were excluded from all other kinematic analyses, and from all EEG analyses.

### Kinematic analysis

The stylus’ position on the tablet was upsampled to 300 Hz and differentiated over time to compute speed. Onset of the centre-out movement was defined as the first time when the stylus’ speed exceeded 2 cm/sec while the red dot was still within 0.45° visual angle (1.6° for Experiment 2) of the screen centre. We determined maximum movement speed between movement onset and 100 ms after crossing the circular array of targets. We then defined movement offset as the sample before speed first fell below 2 cm/sec, following maximum speed. Movement onset and offset were used in computing movement duration, movement distance, and movement curvature (linearity index). The latter was defined as the maximum perpendicular distance of the actual movement trajectory to a straight line connection between the location at movement onset, and the location at movement offset, divided by the distance between home position and endpoint (Atkeson & Hollerbach, 1985). Movement direction was computed as the angle between the point at which the red dot crossed the circular array of targets, and the centre of the home position. By subtracting the direction of the target, we computed target error. Throughout the manuscript, positive target error values correspond to counter-clockwise error.

We excluded trials from kinematic and EEG analyses if movement onset was premature (before the target turned green), if the red dot “froze” in place before crossing the circular array of targets (i.e., if outward movement was too slow or too short), if speed fell below 2 cm/sec before movement offset (i.e., if a movement had more than a single, distinct velocity peak), or if no movement offset could be determined because of ongoing movement up to the point at which the return circle appeared. In addition, some participants reported post-hoc that they forgot to use the aiming strategy in few trials of the rotation block. Consequently, trials in which the target error differed by at least +70° from the mean target error across the entire rotation block were also excluded. In total, these criteria resulted in exclusion of 7.5% of trials in Experiment 1, and of 23.1% of trials in Experiment 2. The relatively high number of excluded trials in Experiment 2 was largely due to frequent premature movement onset in these older-age participants (10% of trials would have been excluded in Experiment 2 had we ignored premature movement onset).

Statistical analyses of learning were based on non-parametric permutation and resampling tests. In Experiment 1, we compared the difference in mean target error between the first vs. last 50 trials (see below) of the rotation block with a distribution under the null hypothesis of no target error difference, obtained from 10000 unique permutations of the labels “first 50 trials” and “last 50 trials” across subjects. A p-value was obtained as the proportion of permutations for which the difference in target error was larger than the actual difference in the non-permuted data. In Experiment 2, we were interested in differences in learning rates between patients and controls. To this end, we extracted learning rate from an established computational model of visuomotor adaptation in the context of instructed strategy use (Taylor & Ivry, 2011). In this model, target error in trial n 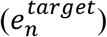 is the difference between the actual visuomotor rotation (r, which is identical across trials) and the aiming strategy (*s*_*n*_), plus the estimated visuomotor rotation in trial n, (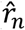; all in degrees):

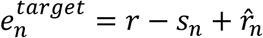

At the beginning of the rotation block,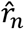 is zero. That is, irrespective of the instructed strategy, which fully compensates for the actual visuomotor rotation, there is a sensory prediction error,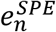,

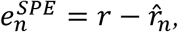

which drives (implicit) adjustments to the estimated visuomotor rotation in the next trial n+1:

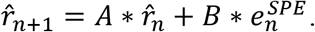

Here, A and B are free parameters representing a memory term and a learning rate, respectively. This simple extension of classic state-space models can reliably reproduce the over-compensation observed in human behavioral data when an instructed strategy is used to compensate for a visuomotor rotation (Mazzoni & Krakauer, 2006; Taylor & Ivry, 2011). Unlike in Taylor and Ivry’s original model (2011), *s*_*n*_ was identical across trials (from the second trial of the rotation block onwards), given that we instructed participants not to deviate from the original strategy.

We estimated A and B separately for patients and controls, using fminsearch.m, and lower/upper bounds for A and B of [0 1] and [-Inf Inf], respectively. To statistically compare learning rates across groups, we randomly reconstructed groups by resampling without replacement 10000 times from the entire cohort of participants in Experiment 2 (patients and controls together). We then computed a p-value as the proportion of resampling iterations for which group differences in learning rate were larger than the actual learning rate difference between the original groups prior to resampling.

### EEG recording and analysis

EEG was recorded using Brain Products amplifiers (Brain Products Inc., Gilching, Germany). The montage included 33 AgCl electrodes referenced against the right mastoid, including a left mastoid electrode, one electrode at each outer canthus, and one electrode below the right eye. Scalp electrodes were arranged in a standard 10 – 20 montage. Impedance was maintained at or below 5 kΩ. Data were sampled at 500 Hz, online low-pass filtered at 250 Hz and online high-pass filtered at .1 Hz.

EEG analysis was conducted in MatLab (The Mathworks Inc., Nattick, Massachusetts, USA) using FieldTrip toolbox (Oostenveld et al., 2011). The EEG was offline band-pass filtered between .1 and 100 Hz (fourth-order Butterworth filter, two-pass), and line noise was removed with a two-pass, fourth-order Butterworth band-stop filter. The EEG data were then epoched from 1500 ms before first target presentation (in white) to the return to the home position, and locked to movement offset. Using ft_rejectvisual.m, epochs were visually inspected, and trials (settings: ‘maxzval’ and ‘meanzval’) or channels (setting: ‘var’) with exceptionally high EEG signal fluctuations were rejected. Eye movement artefacts, including blinks, and ECG artefacts were removed using independent component analysis, followed by a second step of ft_rejectvisual.m. Rejected channels were replaced by a weighted average of their neighbours (ft_channelrepair.m) before data were re-referenced to a common average across scalp electrodes (Alayrangues, Torrecillos, Jahani, & Malfait, 2019; Tan et al., 2016). Fourier-transformed data segments sampled every 20 ms were then multiplied with a Fourier-transformed Hanning taper (of length 400 ms). Spectral power was computed as the squared absolute of the Fourier transform. Given our primary interest in oscillations in the beta frequency range, we focused on frequencies between 2.5 and 40 Hz, analysed in steps of 2.5 Hz. We were particularly interested in the amplitude of the post-movement rebound of beta-oscillations. Given that movement typically results in a beta desynchronization (Pfurtscheller et al., 2003), ongoing or new movement following movement offset would have confounded post-movement beta power. Therefore, we excluded trials from EEG analyses for which movement speed exceeded 5 cm/sec in the first second after movement offset.

Our analyses of post-movement beta power proceeded in two steps. In a first step, we identified the frequency range, time window, and EEG channels that showed a prominent post-movement beta rebound across all four groups of participants, and across all trials. To this end, we constructed group-level time-frequency maps and sensor-level topographies of t-values that compared, across all trials, peri-movement time-frequency modulations to baseline power. As a baseline, we calculated mean power, separately for each frequency bin and EEG channel, across the entire duration of each trial excluding the return movement, i.e., from 1500 ms before first target presentation to the onset of the return circle. Across groups, a cluster of ten channels (including PO3, P3, Pz, CP1, CP2, Cz, C3, FC1, FC2 and Fz), a time window from 350 to 1300 ms after movement offset (see also Kilavik et al., 2013), and a frequency window from 15 to 32.5 Hz captured the post-movement beta rebound (**Fig. 3A** and **5A**). Note that this contrast for identifying channels, time bins, and frequency bins of interest was independent of the contrast in the second-step analysis. In this second step, we compared the relative change in power, compared to baseline power as defined above, across the time-frequency window and channels identified in the first step between different stages of the learning paradigm, as well as between groups. Specifically, we asked whether a decrease of the post-movement beta rebound at the beginning of adaptation, compared to baseline, and an increase via its end (Tan et al., 2016) depends on strategy learning (group A vs. B in Experiment 1), and/or on intact implicit learning from sensory prediction error (patients vs. controls in Experiment 2). To this end, we compared the post-movement beta rebound between the baseline block and the rotation block, where the latter was divided into early and late adaptation trials (first and last 50 trials of the rotation block, respectively). Because group B in Experiment 1, and both groups in Experiment 2, received instructions how to counter the visuomotor rotation only after completing the first two rotation trials, these trials were excluded from analysis in all groups.

**Figure 3.**
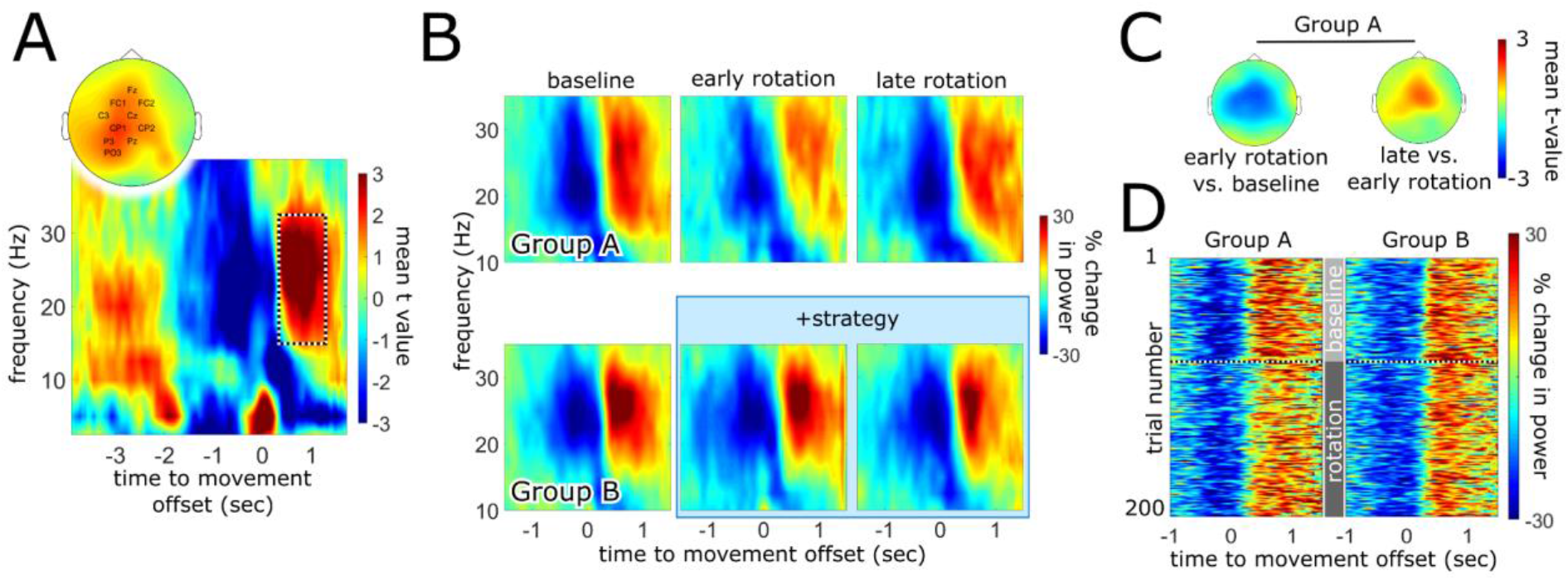
Dynamics of post-movement beta rebound across learning in Experiment 1. A, Time window, frequency range, and channels identified as a time-frequency-channel region of interest (outlined box in time-frequency plot, and channel names) across all trials and groups A and B. Color codes for t-values comparing time-frequency data to baseline power, i.e., power averaged across time, separately for each frequency bin and channel. Zero on the time axes represents movement offset. B, Relative change in peri-movement power (relative to baseline power) in the baseline block (left column), early during adaptation (middle column), and late during adaptation (right column) in groups A (top row) and B (bottom row), averaged across the channels highlighted in panel A. C, Topographies of t-values from a cluster-based permutation test comparing baseline and early rotation (left), and early vs. late rotation (right), in group A. Color represents t-values averaged across a time-frequency cluster within the time-frequency window of interest (panel A). D, Time course of relative change in peri-movement power (relative to baseline power) across all trials in the baseline and rotation blocks (increasing trial order from top to bottom) in groups A (left) and B (right), averaged across the channels highlighted in panel A.

Statistical inference was based on a standard nonparametric permutation test implemented in FieldTrip, which corrects for multiple comparisons across time bins, frequency bins, and channels (Maris & Oostenveld, 2007; details in **Supplementary Material**). Clustering was based on dependent- and independent-samples t-tests across the time window, frequency range, and channels described above. Each test involved 5000 permutations.

To control for effects of movement onset time, duration, maximum speed, curvature, and distance on the dynamics of the post-movement beta rebound, their (linear) effects were estimated based on a multiple linear regression and subtracted from time-frequency data in separate control analyses (ft_regressconfounds; **Supplementary Material**).

## Results

### Experiment 1 – Strategy Learning

Experiment 1 isolated effects of strategy learning over and above any effects of learning from sensory prediction error. To this end, we compared two groups of healthy, young participants, one of which had to discover an aiming strategy (group A), while strategy learning was rendered unnecessary in the other by instructing a fully compensatory strategy (group B).

#### Kinematics

In group A, strategy learning and learning from sensory prediction error were expected to jointly decrease target error across the rotation block. In group B, task instructions obviated strategy learning but not learning from sensory prediction error. Following Mazzoni and Krakauer (2006), persistent sensory prediction error in group B was expected to drive a gradual overcompensation, i.e., an increase in target error opposite to the visuomotor rotation, overriding the instructed strategy.

As expected, participants in both groups made a counter-clockwise target error of approximately 40-45° when first encountering the 45° counter-clockwise visuomotor rotation (**Fig. 2**). In group A, target error then decreased from the first to the last 50 trials of the rotation block (p<0.0001, permutation test; from 17.9 ± 12.6° during early rotation to 6.9 ± 11.1 ° during late rotation; mean ± SD across 50 trials each; **Fig. 2, left**). Despite use of an appropriate aiming strategy from the third trial onwards, which instantly eliminated counter-clockwise target error, group B, too, showed systematic changes of target error across the rotation block, in line with Mazzoni and Krakauer (2006). Specifically, individuals in group B gradually over-compensated throughout the rotation block, producing an increasingly clockwise target error (p<.001 permutation test; from −11.9 ± 6.6° during early rotation to −19.1 ± 7.5° during late rotation (mean ± SD); **Fig. 2, right**). Both groups showed strong after-effects during washout.

Time courses of movement onset time, maximum speed, movement duration, movement distance, and movement curvature for both groups are shown in **Supplementary Fig. 1**.

#### Post-Movement Beta Rebound

Across all blocks, and both groups, we identified a time window, frequency range, and cluster of channels that captured the post-movement beta rebound (**Fig. 3A**). We then asked how the post-movement beta rebound changes from baseline to early adaptation, and from early to late adaptation, when adaptation involves both learning from sensory prediction error as well as strategy learning (group A), and when strategy learning, but not learning from sensory prediction error, is obviated by task instruction (group B).

At baseline, groups A and B had a similar post-movement beta rebound (p>.5, cluster-based permutation test; +14.9 ± 8.4% in group A vs. +15.2 ± 9.8% in group B; mean ± SD; **Fig. 3B**, left). Importantly, however, changes in post-movement beta rebound from baseline to early adaptation differed between the two groups (p=.049 for the interaction of group (A vs. B) and block (baseline vs. rotation), cluster-based permutation test). Specifically, the post-movement beta rebound decreased significantly from baseline to early adaptation in group A (p<.001, cluster-based permutation test; decrease to +9.2 ± 9.3% at early adaptation; mean ± SD; **Fig. 3B**, middle top), but not in group B (p=.44, cluster-based permutation test; +14.1 ± 11.1% at early adaptation; mean ± SD; **Fig. 3B**, middle bottom). As a result, early during adaptation, the post-movement beta rebound tended to be smaller in group A compared to group B (p=.09, cluster-based permutation test).

In addition, groups differed with respect to changes in post-movement beta rebound from early to late adaptation (p=.03 for the interaction of group (A vs. B) and block (early vs. late adaptation), cluster-based permutation test). In group A, there was a significant increase in post-movement beta rebound from early to late adaptation (p=.02, cluster-based permutation test; increase to +13.7 ± 6% at late adaptation; mean ± SD; **Fig. 3B**, right top), which was absent in group B (p>.9, cluster-based permutation test; +12.9 ± 8% at late adaptation; mean ± SD; **Fig. 3B**, right bottom). Taken together, the post-movement beta rebound at central channels (**Fig. 3C**) decreased when an aiming strategy had to be learnt to counter a visuomotor rotation, and then recovered throughout learning, while learning from sensory prediction error alone did not change post-movement beta rebound significantly (**Fig. 3D**).

To ensure that these dynamics across learning stages, and their differences between groups, were not explained by differences in kinematics, we repeated the above analyses after controlling for movement onset time, movement duration, movement speed, movement curvature, and movement distance (**Supplementary Material**).

### Experiment 2 – Learning from Sensory Prediction Error

Experiment 2 examined the dynamics of post-movement beta rebound across visuomotor adaptation in patients with cerebellar ataxia, who have deficits both in learning from sensory prediction error and in strategy learning. To eliminate effects of strategy learning, and isolate effects of sensory prediction error, both patients and age-matched healthy controls deployed an instructed, fully compensatory strategy.

#### Kinematics

Both patients and controls showed a gradual over-compensation across the rotation block (**Fig. 4**), however, with different learning rates. Patients had a significantly lower learning rate than controls (p=.02, resampling test; .005 (patients) vs. .026 (controls)). In particular, target error increased less in the clockwise direction from baseline to the first 50 trials of the rotation block in patients than in healthy controls (p<.01, permutation test; from 2.3 ± 8.9° (baseline, mean ± SD) to −4.7 ± 9.6° (early rotation) in patients, compared to a clockwise increase from 4.2 ± 3.4° (baseline) to −11.1 ± 4.8° (early rotation) in healthy individuals). There was no significant group difference in the memory parameter of the setpoint state-space model (p>.1, resampling test; .98 (patients) vs. .95 (controls)).

**Figure 4.**
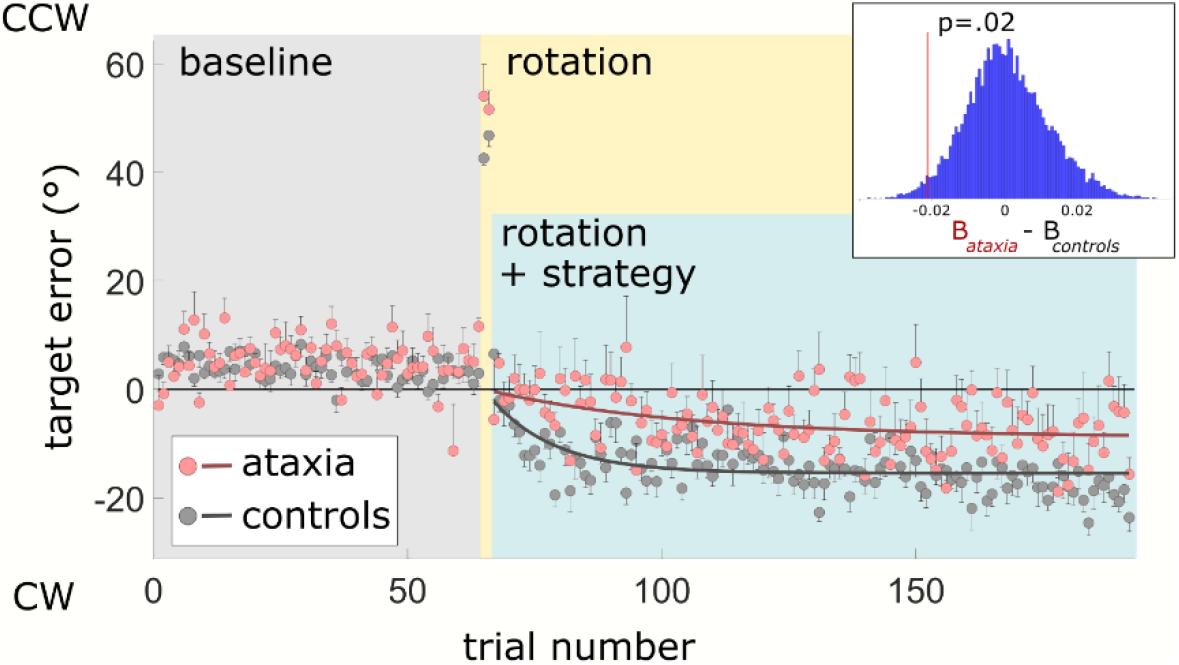
Learning in Experiment 2. Target error as a funcion of experimental block (baseline, rotation + strategy) in patients with cerebellar ataxia (red dots) and age-matched healthy controls (grey dots). Grey and red dots represent the mean, and error bars the standard error of the mean, across participants in each group. The red and grey lines represent learning curves for patients and controls, respectively, predicted by the setpoint state-space model based on group-level target error of the patient and control cohort. The inset shows the actual group difference in learning rate compared to a distribution of group differences under the null hypothesis of no group differences, obtained via resampling without replacement. CCW, counter-clockwise; CW, clockwise.

Time courses of movement onset time, movement speed, movement duration, movement distance, and movement curvature for patients and controls are shown in **Supplementary Fig. 2**. Comparing movement curvature (linearity index) between catch trials and regular trials provided evidence that there was little, if any, feedback correction of movements (**Supplementary Fig. 3**).

#### Post-Movement Beta Rebound

The same time window, frequency range, and cluster of channels as in Experiment 1 also captured the post-movement beta rebound in Experiment 2 (**Fig. 5A**). At baseline, patients tended to have a lower post-movement beta rebound than controls (p=.11, cluster-based permutation test; +4.3 ± 9% in patients vs. +11.5 ± 15.7% in controls; mean ± SD; **Fig. 5B**, left). Importantly, changes in post-movement beta rebound from baseline to early adaptation differed between the two groups (p<.01 for the interaction of group (patients vs. controls) and block (baseline vs. rotation), cluster-based permutation test). Specifically, the post-movement beta rebound increased significantly from baseline to early adaptation in patients (p<.01, cluster-based permutation test; increase to +12.8 ± 16.5% at early adaptation; mean ± SD; **Fig. 5B**, middle top), but not in controls (p>.6, cluster-based permutation test; +13.5 ± 14.5% at early adaptation; mean ± SD; **Fig. 5B**, middle bottom). There was no significant difference between groups with respect to changes in post-movement beta rebound from early to late adaptation (p>.5 for the interaction of group (patients, controls) and block (early vs. late adaptation), cluster-based permutation test). Taken together, the post-movement beta rebound at central and parietal channels (**Fig. 5C**) increased in patients, but not in healthy controls, when an instructed aiming strategy was used to counter a visuomotor rotation (**Fig. 5D**).

**Figure 5.**
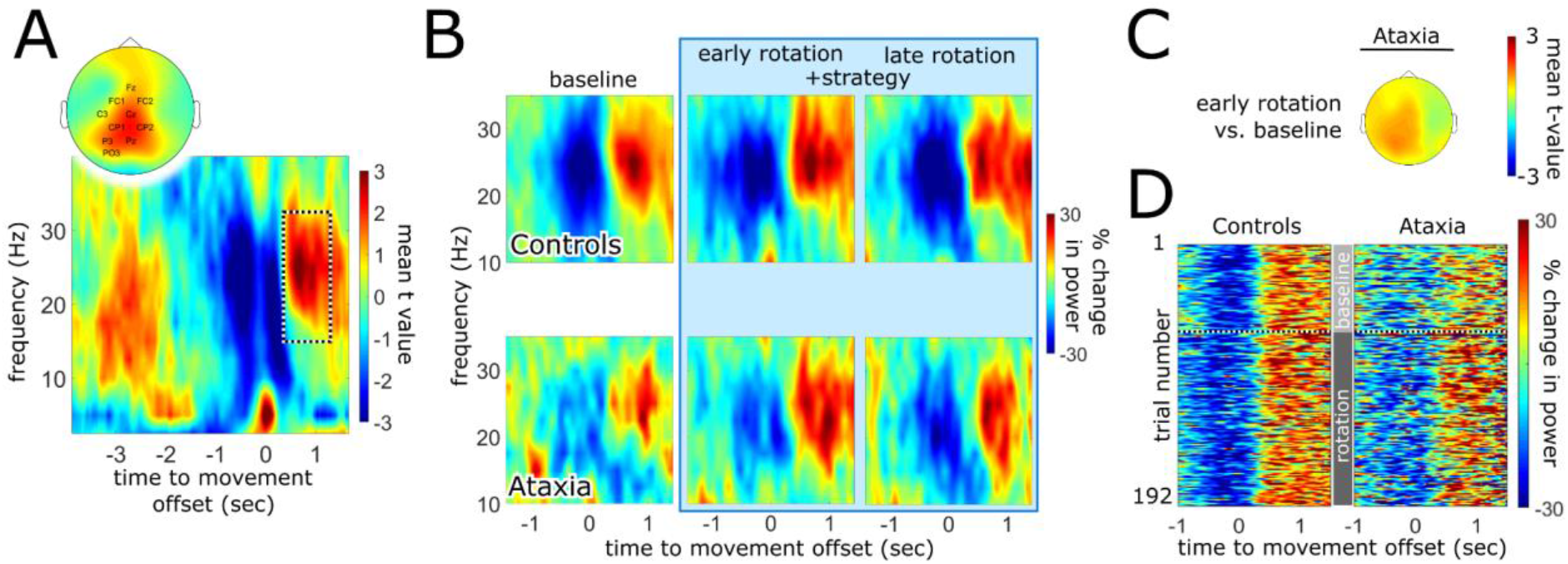
Dynamics of post-movement beta rebound across learning in Experiment 2. A, Time window, frequency range, and channels identified as a time-frequency-channel region of interest (outlined box in time-frequency plot, and channel names) across all trials, as well as patients and controls (identical to Experiment 1). Color codes for t-values comparing time-frequency data to baseline power, i.e., power averaged across time, separately for each frequency bin and channel. Zero on the time axes represents movement offset. B, Relative change in peri-movement power (relative to baseline power) in the baseline block (left column), early during adaptation (middle column), and late during adaptation (right column) in healthy age-matched controls (top row) and patients with cerebellar ataxia (bottom row), averaged across the channels highlighted in panel A. C, Topography of t-values from a cluster-based permutation test comparing baseline and early adaptation in patients. Color represents t-values averaged across a time-frequency cluster within the time-frequency window of interest (panel A). C, Time course of relative change in peri-movement power (relative to baseline power) across all trials in the baseline and rotation plus strategy blocks (increasing trial order from top to bottom) in age-matched controls (left) and patients (right), averaged across the channels highlighted in panel A.

To ensure that these dynamics across learning stages, and their differences between groups, were not explained by differences in kinematics, we repeated the above analyses after controlling for movement onset time, movement duration, movement speed, movement curvature, and movement distance (**Supplementary Material**).

## Discussion

Cerebellar ataxia is increasingly recognised as a disorder that affects several interacting motor learning mechanisms (Therrien et al., 2016; Butcher et al., 2017; Donchin and Timmann, 2019; Wong et al., 2019). Understanding these interactions, and in particular their relevance to the pathophysiology of cerebellar ataxia, likely requires a network perspective that includes extra-cerebellar regions, in particular cerebral cortex. While previous studies have established links between motor learning and cortical beta-oscillations (Alayrangues et al., 2019; Darch, Cerminara, Gilchrist, & Apps, 2020; Espenhahn et al., 2019; Haar & Faisal, 2020; Jahani, Schwey, Bernier, & Malfait, 2020; Palmer et al., 2019; Pollok, Latz, Krause, Butz, & Schnitzler, 2014; Tan et al., 2016; Tan et al., 2014; Torrecillos et al., 2015), the involvement of cortical beta-oscillations in specific learning mechanisms has remained largely unclear, leave alone their relevance to motor learning dysfunction in cerebellar ataxia.

Our study advances the field in several ways. First, it differentiates between learning a strategy, and learning from sensory prediction error, and demonstrates that strategy learning decreases the post-movement beta rebound, over and above any influence of learning from sensory prediction error. However, our study also demonstrates that explicit, strategy-based control, and implicit control guided by sensory prediction error, share as a common neural pathway a modulation of the post-movement beta rebound. Specifically, a dysfunction of implicit, sensory prediction error-driven learning in cerebellar ataxia results in an increase of the post-movement beta rebound once an explicit strategy is used to counter a visuomotor rotation. We propose that this common pathway may represent a physiological correlate of an interaction between strategy learning and learning from sensory prediction error proposed in the literature (Donchin and Timmann, 2019; Wong et al., 2019). Specifically, the increase in post-movement beta rebound we observe in patients during strategy use (Experiment 2), together with our finding that adjustments to a strategy normally decrease the post-movement beta rebound (Experiment 1), may explain why patients fail to develop sufficient strategies on their own (Vaca-Palomares et al., 2013; Butcher et al., 2017).

### Post-Movement Beta Rebound and Motor Learning

Tan et al. (2014) first reported that the rebound of cortical beta-power which has long been known to follow movements (Pfurtscheller, 1992) depends on a combination of movement error and error history. In a follow-up study that further emphasised the role of context for this modulation (Tan et al., 2016), they concluded that the post-movement beta rebound indexes confidence in a current internal model. Other groups have since confirmed that movement error results in lower post-movement beta rebound (Alayrangues et al., 2019; Palmer et al., 2019; Torrecillos et al., 2015). Based on these findings, and the role of movement error in influential computational theories of motor control (Wolpert and Miall, 1996; Körding and Wolpert, 2004; Shadmehr et al., 2010), it has been proposed that peri-movement cortical beta-oscillations are involved in dynamically adjusting (confidence in) internal models that enable movement execution and motor learning (Cao & Hu, 2016; Palmer et al., 2019; Palmer et al., 2016; Tan et al., 2016).

These ideas are broadly compatible with theoretical frameworks of beta-oscillations as promoters of a “status quo” (Engel and Fries, 2010) that favors incumbent over novel processing (Brittain and Brown, 2014), possibly by constraining information coding capacity through neural synchrony (Baker, Kilner, Pinches, & Lemon, 1999; Brittain & Brown, 2014; Zohary, Shadlen, & Newsome, 1994; see also Hosaka et al., 2016). However, the design of previous studies into learning-related dynamics of the post-movement beta rebound complicates their interpretation with respect to such specific learning mechanisms as internal model adaptation. By design, these studies cannot dissociate learning from sensory prediction error and learning a cognitive strategy. Here, by obviating strategy learning in one group of young, healthy participants, but not the other, we isolate effects of strategy learning over and above learning from sensory prediction error.

What specific aspect of strategy learning could be responsible for the observed decrease in post-movement beta rebound? Previous studies have proposed that the dynamics of the post-movement beta rebound during learning reflect movement error, i.e., a discrepancy between desired and actual movement (“target error” in Figures 2 and 4, as well as in Taylor & Ivry, 2011, and in Taylor et al., 2014), even when that error does not result in adjustments of behavior (Torrecillos et al., 2015). Target error is considered a critical feedback for learning an aiming strategy (Taylor & Ivry, 2011; Taylor et al., 2014). In Experiment 1, participants in group A, who showed a decrease in post-movement beta rebound early during adaptation, also produced, on average, larger target errors at this stage, compared to group B (Figure 2). However, two aspects of our results speak against an exclusive function of the post-movement beta rebound as a generic error signal. First, given that target error subsequently increased during adaptation in group B, a generic error signal should change in amplitude from early to late adaptation in that group, even when that error does not result in behavioral (strategy) adjustments (Torrecillos et al., 2015). However, there was no significant change in the amplitude of the post-movement beta rebound from early to late adaptation in group B (Figure 3). Consequently, if the dynamics of the post-movement beta rebound indeed reflects target error, then it does so likely in a context of its behavioral relevance (Tan et al., 2014) or salience (Alayrangues et al., 2019). Second, and more importantly, the increase in post-movement beta rebound from baseline to early adaptation observed in patients (Figure 5) is difficult to reconcile with an exclusive interpretation of the post-movement beta rebound as an error signal.

### Post-Movement Beta Rebound During Learning in Ataxia

This increase in post-movement beta rebound from baseline to early adaptation in patients coincided with two concurrent changes in their behavioral task. At the beginning of the rotation block, visuomotor contingencies changed, causing a sensory prediction error. At the same time, patients started using, and maintaining, an instructed strategy to counter the change in visuomotor contingencies (with a delay of two trials excluded from analysis). Was the observed increase in post-movement beta rebound in patients driven by the change in visuomotor contingency, the use/maintenance of a strategy, or a combination of both? Following ideas that beta-signals in EEG reflect superposition of several functional processes (Kilavik et al., 2013), we think the most plausible and parsimonious explanation at this stage, which accounts for results of Experiments 1 and 2, is a combination of how patients encode, or use, sensory prediction error, and a physiological effect of maintaining a strategy, as follows.

Our behavioral results demonstrate reduced adaptation of an internal visuomotor model by sensory prediction error in patients (Figure 4), in line with Taylor et al. (2010). Impaired adaptation of internal models in response to sensory prediction error is considered a key learning dysfunction in cerebellar ataxia (Smith and Shadmehr, 2005; Tseng et al., 2007; Synofzik et al., 2008; Izawa et al., 2012). This dysfunction is thought to exist independently of any experimentally induced changes in sensorimotor contingencies (Bhanpuri et al., 2014). Accordingly, we would expect miscalibrated internal models in patients already at baseline, before the visuomotor rotation is introduced, given that the visuomotor mapping at baseline was itself a newly learnt, anisotropic mapping of stylus movements on the tablet (in the horizontal plane) to feedback on the monitor (in the vertical plane). Any EEG signal modulation that reflects this miscalibration, or the ensuing sensory prediction error, should therefore be in the same direction at baseline and early during adaptation. However, at baseline, the post-movement beta rebound tended to be attenuated in patients, compared to controls (see also Aoh et al., 2019), while it increased from baseline to early adaptation in patients. This pattern seems difficult to reconcile with the idea that the observed changes in the dynamics of the post-movement beta rebound in patients purely reflect abnormal encoding, or use, of sensory prediction error.

Instead, we think the increase in post-movement beta rebound from baseline to early adaptation reflects an effect of strategy use, or strategy maintenance, which, in patients, is not balanced by an opposing effect of sensory prediction error that is present in healthy individuals (or an effect of sensory prediction error-driven internal model adaptation). As discussed above, cortical beta-oscillations have been associated with maintenance of a current “motor set” (Engel and Fries, 2010), e.g., with steady-state force output or maintaining a posture (Baker et al., 1999; Kilner et al., 2000; Gilbertson et al., 2005; Schoffelen et al., 2005). Hosaka et al. (2016) showed that beta-power increased in the supplementary and pre-supplementary motor areas (before a Go cue) when monkeys were producing a learnt sequence of arm movements from memory, compared to trials that required updating that sequence based on new instructions (see also Jahani et al., 2020).

We think that using, or maintaining, an (instructed) strategy in our task may similarly increase the post-movement beta rebound, and that sensory prediction error, or the resulting adaptation of an internal model, may decrease the post-movement beta rebound. This could explain observed patterns across Experiments 1 and 2 (**Fig. 6**). An increase in post-movement beta rebound upon strategy maintenance (magenta box), would prevail when an opposing decrease by sensory prediction error, or internal model adaptation, is reduced due to impaired implicit learning (yellow box), such as observed in cerebellar ataxia (yellow/magenta box). In healthy individuals, however, the increase due to using, or maintaining, a strategy (magenta box) would be balanced by a strong decrease due to sensory prediction error, or internal model adaptation (blue box), resulting in no net change, as in group B in Experiment 1, and in controls in Experiment 2 (blue/magenta box). Learning a strategy, or target error (green box), and (learning from) sensory prediction error (blue box), on the other hand, would jointly decrease the post-movement beta rebound, as in group A in Experiment 1 (blue/green box). Finally, the latent miscalibration of internal models in cerebellar ataxia (Bhanpuri et al., 2014) would result in a (slight) decrease of the post-movement beta rebound at baseline (yellow box, and Aoh et al., 2019).

**Figure 6.**
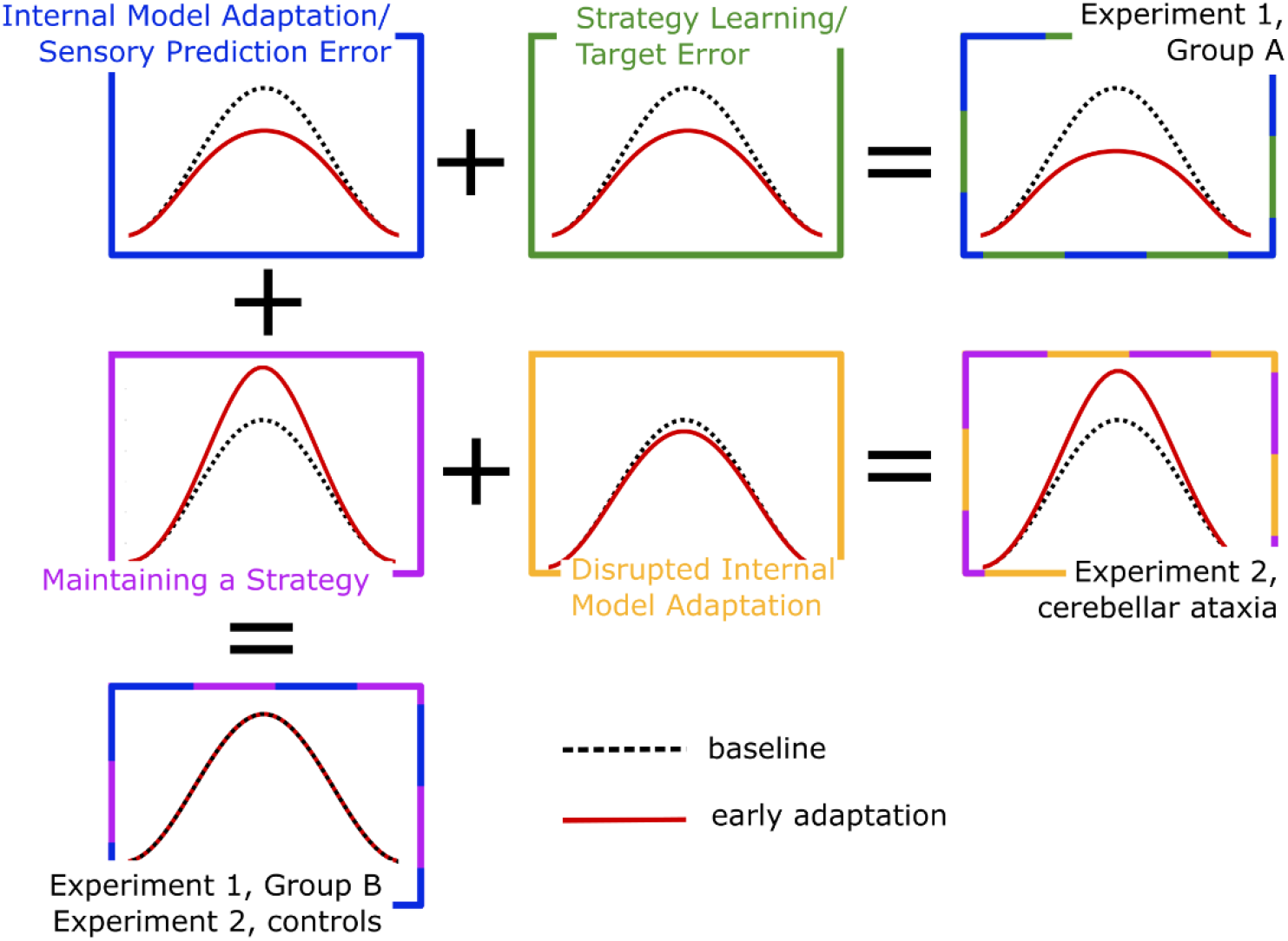
A model of learning-related dynamics of the cortical post-movement beta rebound. Understanding learning-related dynamics of cortical beta-oscillations may have strong implications for other disorders. Similarly to patients with cerebellar ataxia (Figure 5B, left, and Aoh et al., 2019), the post-movement beta rebound is decreased in patients with Parkinson’s Disease (Pfurtscheller et al., 1998; Degardin et al., 2009), and in patients with obsessive-compulsive disorder (Leocani et al., 2001). Interestingly, patients with Parkinson’s Disease display impaired learning specifically when large visuomotor perturbations are introduced abruptly (Contreras-Vidal and Buch, 2003; Mongeon et al., 2013), which is thought to favour explicit strategy use (Morehead et al., 2015). Obsessive-compulsive disorder, on the other hand, has been associated with altered function of internal forward models (Gentsch et al., 2012).

The net increase in post-movement beta rebound when patients start using an aiming strategy to counter a visuomotor rotation, together with the decrease of this signal normally observed during strategy learning in healthy individuals, could explain why patients are less flexible in adjusting strategies, e.g., why they develop only incomplete strategies (figure 2B in Butcher et al., 2017). At this stage, however, the above account remains somewhat speculative, in particular because it is difficult to fully isolate learning a strategy from the change in visuomotor contingencies it aims to compensate for, and, thus, from sensory prediction error.

### Limitations

The patient cohort studied here was heterogeneous with respect to specific etiologies of cerebellar degeneration, as well as regarding disease stage. Disease stage, in particular, may influence to what extent interactions between motor learning mechanisms play a role in cerebellar ataxia (Donchin and Timmann, 2019). The heterogeneity in etiology may explain why patients showed only a trend for attenuated post-movement beta rebound at baseline, compared to controls. In a more homogeneous cohort of patients with spinocerebellar ataxia type 3, Aoh et al. (2019) have recently reported a significant decrease in post-movement beta rebound following self-paced wrist extensions.

We propose that modulations of the post-movement beta rebound both by strategy learning (Experiment 1), and by learning from sensory prediction error (Experiment 2), represent a candidate physiological interaction between these learning mechanisms. However, the observed modulations of the post-movement beta rebound in EEG may map onto several, functionally and/or spatially distinct populations of neurons. We cannot exclude the possibility that recording techniques with higher spatial resolution than achieved with EEG would dissociate neurons responsible for a modulation of post-movement beta-oscillations for strategy learning from neurons involved in learning from sensory prediction error. Single-and multi-unit recordings in non-human primates may address this issue in the future.

Finally, it has been shown that the post-movement beta rebound varies with post-movement electromyographic activity (Demandt et al., 2012; Kilavik et al., 2013). While we excluded trials from EEG analysis based on a post-movement velocity threshold, we cannot exclude the possibility that ongoing electromyographic activity, even with no, or little, overt movement, varied systematically with learning in our experiments (Osu et al., 2002). While several groups have proposed that training to modulate end-point stiffness could benefit patients with cerebellar ataxia (Bastian et al., 1996; Scheidt et al., 2011), Gibo et al. (2013) have provided evidence that cerebellar pathology may actually impair stiffness control. While our own study did not include any measure of muscle co-contraction, we think the pattern of results cannot be easily explained purely by changes in electromyographic activity associated with arm stiffness. Specifically, for changes in arm stiffness to explain the observed pattern of post-movement beta modulations, any difference in terminal co-contraction between patients and controls would have to vanish once patients use an aiming strategy (Fig. 5B, middle vs. left). In addition, Gibo et al.’s finding of diminished impedance control in patients with cerebellar ataxia contrasts with the *reduced* post-movement beta rebound we observed in patients at baseline. Following Demandt et al.’s (2012) findings, such a reduction in post-movement beta rebound would have to be explained by stronger, rather than diminished, (end-point) impedance control.

### Conclusions

Our study extends a recently emerging network perspective on the electrophysiology of cerebellar ataxia that includes cerebral cortex (Lu et al., 2008; Aoh et al., 2019; Visania et al., 2020), and turns to functionally relevant, learning-related modulations of cortical signals. Our results point to a common neural pathway of strategy learning and learning from sensory prediction error, reflected in a modulation of the sensorimotor cortical, post-movement beta rebound. This modulation is out of balance in cerebellar ataxia, an imbalance that could represent a candidate physiological phenomenon of impaired strategy learning in cerebellar ataxia. Cerebellar degeneration thus surfaces in learning-related, peri-movement oscillatory signals at the level of cerebral cortex.

## Acknowledgements

We thank Laura Herrmann for excellent technical support, all participants for their time and effort, and all members of the Motor Learning research group for their valuable feedback on the manuscript.

