## Supplementary Material for "Altered Dynamics of Cortical Beta-Oscillations during Motor Learning in Cerebellar Ataxia"

### Supplementary Methods

*Statistical inference on EEG data*

Statistical inference was based on a standard nonparametric permutation test implemented in FieldTrip, which corrects for multiple comparisons across time bins, frequency bins, and channels (Maris & Oostenveld, 2007; details in **Supplementary Material**). First, a t-statistic (dependent-samples t-test) was computed for a given contrast of interest (e.g., early adaptation vs. baseline) for each time bin, frequency bin, and channel separately. Adjacent time bins and frequency bins and neighbouring channels whose t-statistic exceeded a threshold ($p<0.05$, 2-sided) were then clustered. For each cluster, a cluster-level statistic was obtained by summing t-values across its elements. The null hypothesis was rejected if this cluster-level statistic exceeded a critical value estimated from a distribution obtained by at least 5000 random data permutations. To this end, the two components of a contrast (e.g., early adaptation vs. baseline) were randomly assigned the two labels ("baseline," "early adaptation") independently for each subject. For each permutation, a t-statistic was recomputed for each time/frequency bin in each channel. Adjacent bins whose t-statistic exceeded a two-sided threshold of $p<0.05$ were then clustered, and all t-values in a given cluster were summed to obtain a cluster-level statistic. The maximum (positive and negative) cluster-level statistic from each permutation was used to build a nonparametric distribution, reflecting the null hypothesis of no difference. A p-value was defined for each positive and negative cluster in the actual data as the proportion of random permutations with a higher or lower maximum cluster-level statistic, respectively. Clusters in the actual data for which this proportion exceeded a threshold of $p<0.05$ (two-sided) were considered significant.

*Catch trial analysis in Experiment 2*

To compare movement curvature (linearity index) between catch trials and rotation trials in Experiment 2, we used a Bayesian repeated-measures analysis of variance (ANOVA) in JASP (JASP Team, 2019). Specifically, we compared models that did, or did not, include "trial type" (two levels: catch trial, rotation trial) as a factor in a Bayesian repeated ANOVA. The second factor in this ANOVA was group (patients vs. controls). Linearity index was log-transformed for normality.

### Supplementary Results

*Dynamics of post-movement beta rebound, controlling for kinematics*

Experiment 1

To ensure that dynamics of the post-movement beta rebound across learning stages, and their differences between groups, in Experiment 1 were not explained by differences in kinematics, we repeated the EEG analyses after controlling for movement onset time, movement duration, movement speed, movement curvature, and movement distance (**Supplementary Fig. 1**). At baseline, groups A and B had a similar post-movement beta rebound (p>.6, cluster-based permutation test; +14.9 ± 8.4% in group A vs. +14.6 ± 8.1% in group B; mean ± SD). However, importantly, changes in post-movement beta rebound from baseline to early adaptation differed between the two groups (p=.04 for the interaction of group (patients vs. controls) and block (baseline vs. rotation), cluster-based permutation test). Specifically, the post-movement beta rebound decreased significantly from baseline to early adaptation in group A (p<.001, cluster-based permutation test; +10.1 ± 8.3% at early adaptation; mean ± SD), but not in group B (p=.42, cluster-based permutation test; +14.5 ± 10.9% at early adaptation; mean ± SD). As a result, early during adaptation, the post-movement beta rebound tended to be smaller in group A compared to group B (p=.1, cluster-based permutation test).

In addition, groups differed with respect to changes in post-movement beta rebound from early to late adaptation (p=.047 for the interaction of group (patients, controls) and block (early vs. late adaptation), cluster-based permutation test). In group A, there was a trend for an increase in post-movement beta rebound from early to late adaptation (p=.09, cluster-based permutation test; +14.1 ± 6.2% at late adaptation; mean ± SD), which was absent in group B (p>.9, cluster-based permutation test; +13.5 ± 8.2% at late adaptation; mean ± SD). Taken together, controlling for effects of movement onset time, movement duration, movement speed, movement curvature, and movement distance on post-movement beta rebound did not qualitatively change statistical results in Experiment 1.

Experiment 2

Similarly, to ensure that dynamics of the post-movement beta rebound across learning stages, and their differences between groups in Experiment 2 were not explained by differences in kinematics, we repeated the EEG analyses of Experiment 2 after controlling for movement onset time, movement duration, movement speed, movement curvature, and movement distance (**Supplementary Fig. 2**). At baseline, there was a weak trend for a lower post-movement beta rebound in patients compared to controls (p=.17, cluster-based permutation test; +4.4 ± 9.3% in patients vs. +11.7 ± 15% in controls; mean ± SD). Importantly, changes in post-movement beta rebound from baseline to early adaptation differed between the two groups (p=.03 for the interaction of group (patients vs. controls) and block (baseline vs. rotation), cluster-based permutation test). Specifically, the post-movement beta rebound increased significantly from baseline to early adaptation in patients (p=.02, cluster-based permutation test; increase to +12.5 ± 15.9% at early adaptation; mean ± SD), but not in controls (p>.9, cluster-based permutation test; +13.5 ± 13.5% at early adaptation; mean ± SD). There was no significant difference between groups with respect to changes in post-movement beta rebound from early to late adaptation (p>.35 for the interaction of group (patients, controls) and block (early vs. late adaptation), cluster-based permutation test). Taken together, controlling for effects of movement onset time, movement duration, movement speed, movement curvature, and movement distance on post-movement beta rebound did not qualitatively change statistical results in Experiment 2.

*Feedback corrections in Experiment 2*

Comparing movement curvature (linearity index) between catch trials and regular trials provided evidence that there was little, if any, feedback correction of movements. Specifically, a model including trial type as a factor (two levels: catch trials, rotation trials), besides group (patients vs. controls), was 2.7 times less likely than the best model, including only group (Bayes Factor; see **Supplementary Fig. 3**).

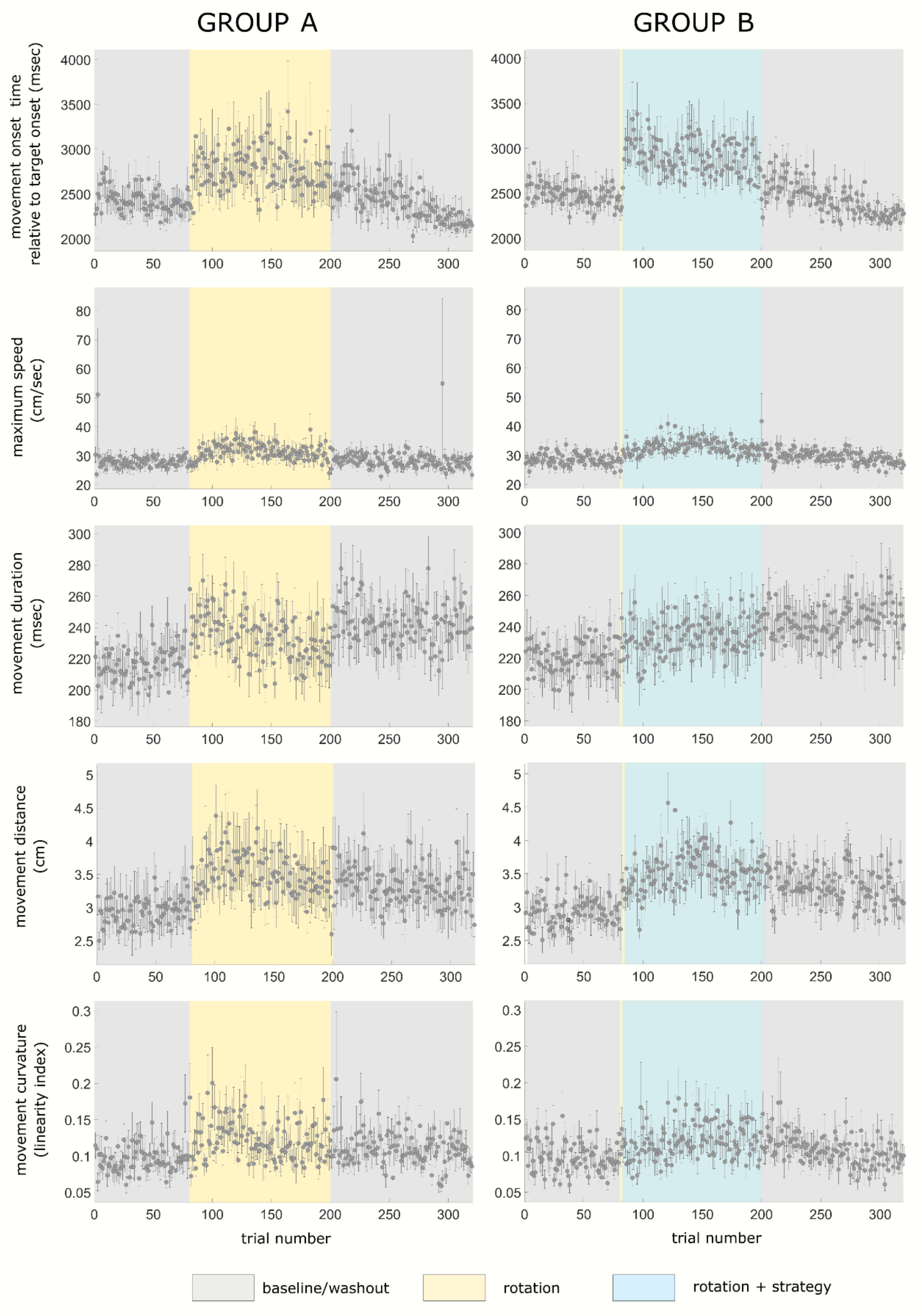


**Supplementary Figure 1. Time courses of movement kinematics across baseline, rotation, and washout blocks in Experiment 1.** Grey dots and error bars represent the mean and standard error of the mean, respectively, across participants in group A (left column) and group B (right column).

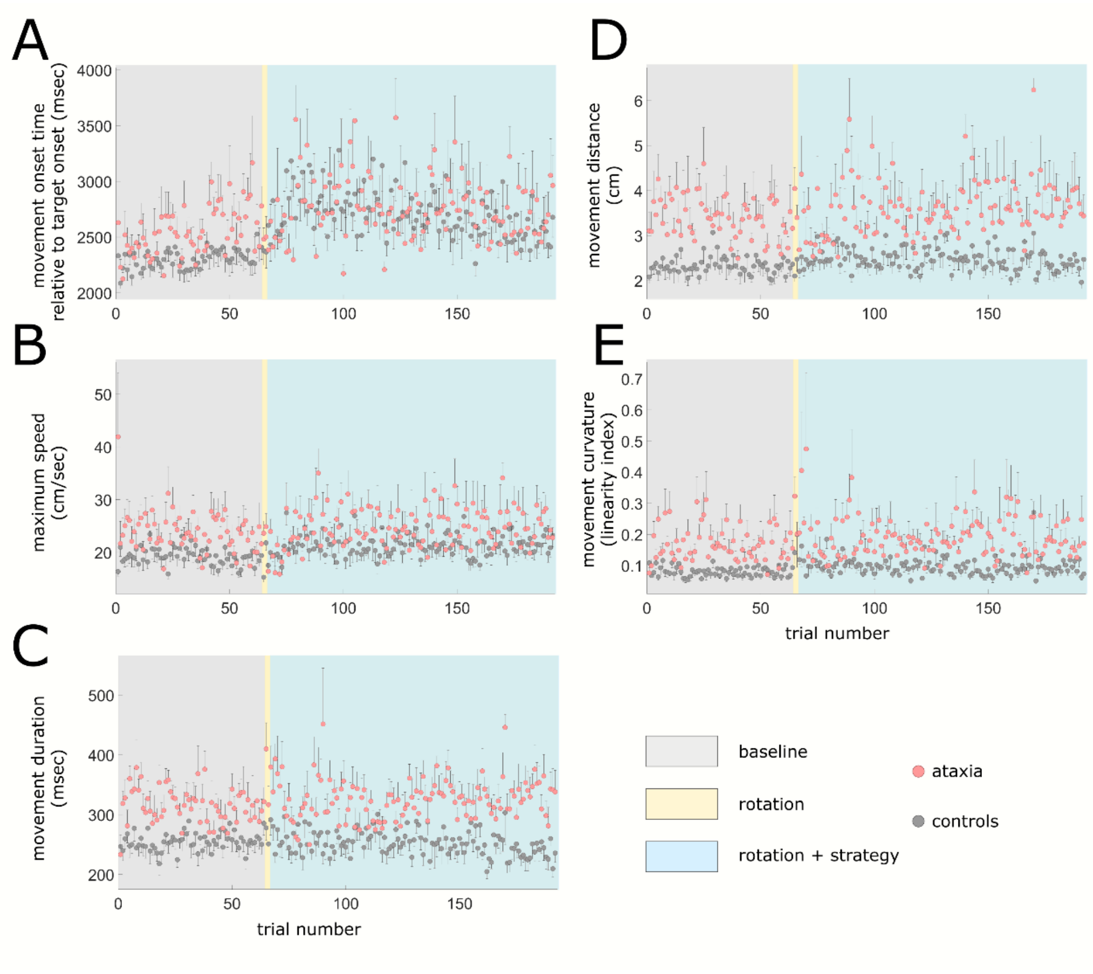


**Supplementary Figure 2. Time courses of movement kinematics across baseline, rotation, and washout blocks in Experiment 2.** Grey and red dots represent the mean, and error bars the standard error of the mean, across patients with cerebellar ataxia (red dots) and age-matched controls (grey dots).

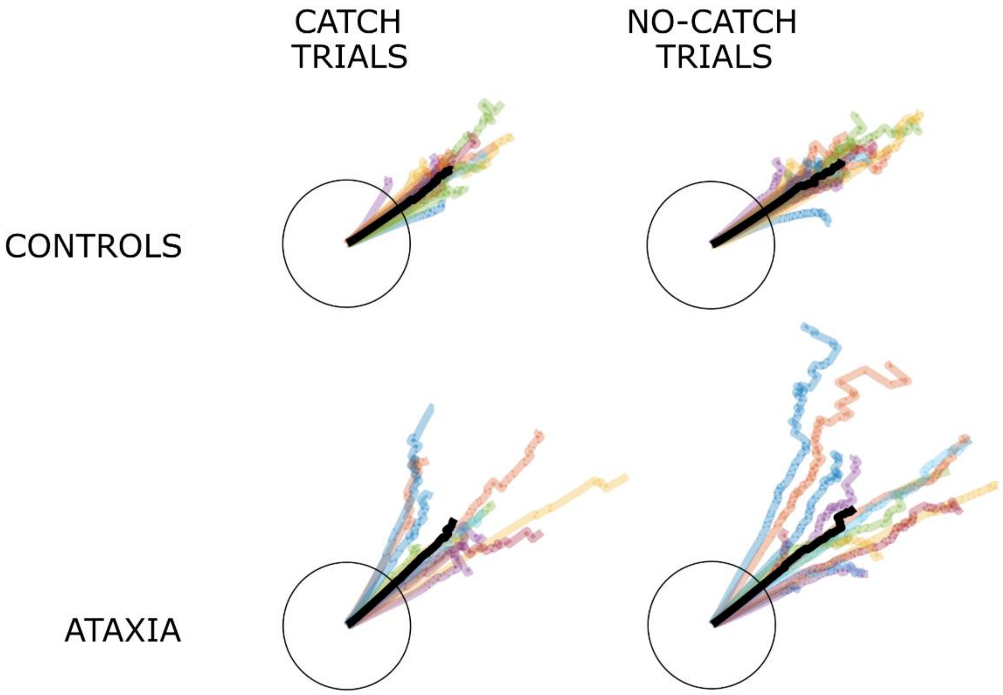

**Supplementary Figure 3. Subject- and group-level average movement trajectories for catch trials (left column) and rotation trials (right column) for patients (lower row) and age-matched controls (upper row).** For illustration purposes, targets are aligned to 12 o'clock noon. Note that patients and controls were deploying an aiming strategy, hence trajectories are directed at the neighbouring target at around 45°. Trajectories in color represent individual subject averages, while black trajectories represent group-level averages. Black circles represent the movement extent required to reach the circular array of targets.
