## Supplementary material for "Altered Dynamics of Cortical Beta-Oscillations during Motor Learning in Cerebellar Ataxia": Table 1

| | Gender | Age | Diagnosis | ICARS Total/100 | ICARS Limb ataxia/52 |
|---|---|---|---|---|---|
| **A001** | M | 75 | Family history, likely dominant inheritance | 34 | 18 |
| **A002** | F | 39 | Family history | 62 | 32 |
| **A003** | M | 49 | SCA2 | 35 | 19 |
| **A004** | F | 33 | Family history, likely dominant inheritance | 22 | 11 |
| **A005** | M | 75 | Family history, likely dominant inheritance | 28 | 14 |
| **A006** | F | 45 | SAOA | 28 | 12 |
| **A007** | F | 63 | SAOA | 22 | 10 |
| **A008** | F | 54 | SCA14 | 32 | 16 |
| **A009** | F | 58 | SCA14 | 33 | 18 |
| **A010** | M | 68 | SCA1 | not available | |
| **A011** | F | 55 | SCA6 | not available | |
| **A012** | M | 67 | Sporadic | 50 | 23 |
| **A013** | M | 52 | SCA4 | 17 | 9 |
| **A014** | F | 60 | SCA1 | 27 | 13 |
| **A015** | M | 69 | Family history, likely dominant inheritance | 30 | 11 |
| **Patients** | F=8 | 57.5 | | | |
| **Controls** | F=7 | 56.4 | | | |

ICARS, International Cooperative Ataxia Rating Scale; SAOA, sporadic adult-onset ataxia; SCA, spinocerebellar ataxia.
